# Reduced efficacy of an anti-toxin vaccine from senescence-driven attenuation of toxin virulence

**DOI:** 10.1101/2025.08.14.670416

**Authors:** Xin Du, Ching-Wen Tseng, Elisabet Bjånes, Hunter Gage, Jaclyn Swan, Chih-Ming Tsai, Irshad Hajam, Cesia Gonzalez, Brian Lin, Victor Nizet, George Y Liu

**Affiliations:** Department of Pediatrics, University of California San Diego, San Diego, California, United States; Division of Infectious Diseases, Rady Children’s Hospital San Diego, San Diego, California, United States; Orca Bio, Menlo Park, California, United States; Department of Pharmacy, University of California San Diego, San Diego, California, United States; Department of Ecological Plant and Animal Sciences, La Trobe University, Melbourne, Australia

**Keywords:** *S. aureus*, vaccine, virulence, immunocompromised, ADAM10, Hla, aged, senescence

## Abstract

It remains unclear why vaccines targeting prominent microbial virulence factors often fail in clinical trials. Because microbial virulence depends on interaction with the host immune system, we investigated how changes in host immune function alter vaccine efficacy. Using a vaccine against *Staphylococcus aureus* alpha toxin (Hla), which targets host metalloprotease ADAM10 on myeloid cells, we show that Hla virulence is reduced in aged mice due to diminished ADAM10 activity and impaired myeloid cell function. Depletion of myeloid cells with cyclophosphamide in young mice similarly reduced toxin virulence. Immunization against Hla conferred strong protection against *Staphylococcus aureus* infection in young but not aged mice. These findings indicate that pathogenic functions of microbial factors characterized in immunocompetent young animals may not predict virulence or vaccine efficacy in immunocompromised hosts. These findings underscore the need to account for host immune status in the development and evaluation of vaccines targeting microbial virulence factors.

## Introduction

Studies of host-pathogen interaction provide the fundamental basis for understanding how microbes establish infection and inflict damage in host tissues. Prominent virulence factors unveiled from these studies serve as targets for translational intervention ^1–3^, providing a rationale for sustained public support of host-pathogen research ^4^. Microbial virulence is most investigated in young adult laboratory rodents at the peak of immune competence. Yet infectious pathology arises from dynamic interactions between microbial factors and host immune components, including activation of downstream effectors and evasion of immune defenses^3^. Thus, the manifestation of microbial virulence depends on an optimally functioning host immune system.

However, immune functions vary substantially across the human lifespan and with clinical conditions such as prematurity, aging, diabetes, or immunosuppressive drug use. These immune-altering states may shift the balance of host-pathogen interactions and thereby affect the expression or impact of microbial virulence. We hypothesized that virulence defined in young adult animals may not translate directly to immunocompromised hosts and that this discrepancy could contribute to the failure of vaccines that target virulence factors.

To test this hypothesis, we focused on alpha-toxin (Hla), a major *Staphylococcus aureus* virulence determinant that has been extensively studied in young rodents ^5,6^. Hla exerts its pathogenic effects through ligation of the host metalloprotease ADAM10, leading to cellular damage, skin barrier disruption, and exacerbated inflammation ^7^. Upon binding ADAM10, Hla triggers cleavage of junctional and adhesion molecules and activates proinflammatory signaling cascades that amplify tissue injury^7–9^. Hla contributes to the pathogenesis of multiple *S. aureus* infections ^5,6^, which collectively accounted for over one million deaths worldwide in 2019 ^10^.

Despite decades of research and over 10 phase 2 or 3 clinical trials, there is still no licensed vaccine against *S. aureus* to date^11,12^. Various explanations have been proposed, including poor immunogenicity, suboptimal adjuvants, and inappropriate trial design that enrolled patients with impactful comorbidities ^13^. A particularly relevant example is the trial of Suvratoxumab, a human monoclonal antibody targeting Hla, which failed to meet its primary endpoints in preventing *S. aureus* pneumonia ^14^. Notably, a subgroup analysis showed that Suvratoxumab was effective in patients under 65 years of age but conferred no benefit in those over 65 ^14^.

This trial outcome raises the critical question of whether age-associated changes in immune function alter the pathogenic potential of virulence factors like Hla. In this study, we tested whether Hla-mediated pathology is reduced in aged or immunosuppressed hosts and whether such alterations influence vaccine efficacy. Our findings have implications for understanding why some virulence-targeting interventions succeed in preclinical models yet fail in real-world clinical settings.

## Results

### Hla virulence is diminished in aged mice

Two well-characterized pathologic functions of Hla are its ability to lyse a broad range of host cells and to induce pro-inflammatory cytokine release. To assess whether senescence impacts Hla pathological functions, we incubated neutrophils purified from young and aged mice or human donors with the toxin. In our study, “young” refers to mice 6-12 weeks of age or humans aged 18-35 years, and “aged” refers to mice aged 16 to 22 months or humans aged 65-85 years.

Neutrophils from aged humans and mice were significantly more resistant to cytolysis than those from young donors (**Figures 1A-B**). IL-1β release was also lower in aged hosts; while differences in mouse IL-1β were modest and did not reach statistical significance, the trend was consistent with human data (**Figure 1C-D**). In vivo, subcutaneous injection of Hla induced significantly smaller dermonecrotic lesions in aged mice than in young mice (**Figures 1E and S1A**). Subcutaneous challenge with wild-type USA300 or an isogenic Hla deletion mutant (Δhla) corroborated these findings: over a 7-day infection, Hla contributed to larger lesion sizes and greater skin bacterial burden in young mice but had a modest effect in aged mice (**Figures 1F-G and S1B**). Notably, *S. aureus* dissemination to deep tissues was greater in aged mice and was independent of Hla activity **(Figure 1H-I)**. These findings provide direct evidence that Hla virulence is attenuated in aged hosts, both in vitro and in vivo, and suggest that host immune status modulates the pathological effects of this well-established toxin.

**Fig. 1.**
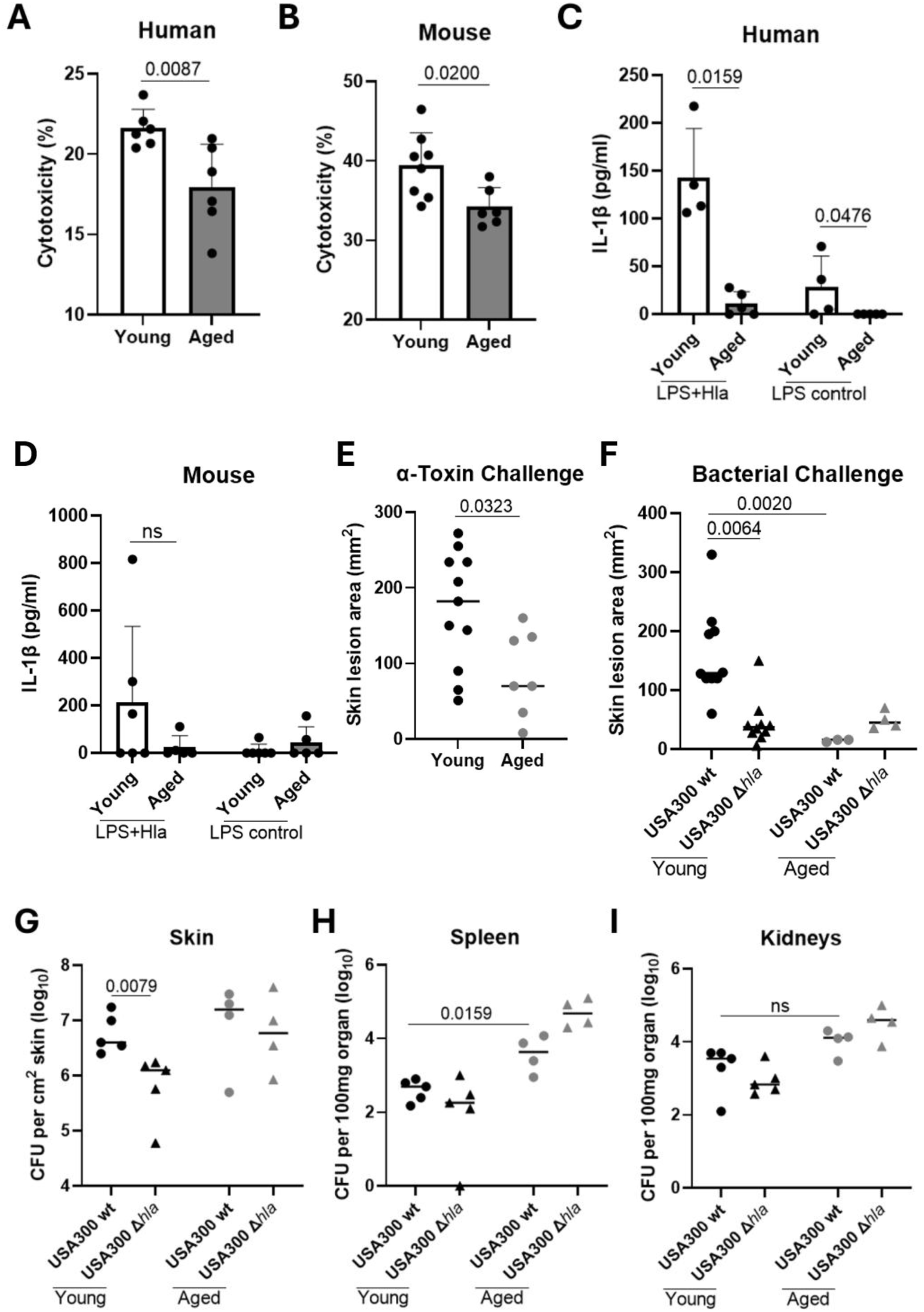
Hla virulence is diminished in aged mice. (A) Hla lysis of human PMNs assessed using LDH assay. Percent cell lysis in baseline untreated control – Young 13.8, aged 14.6 (n=6 humans). 10 U/ml Hla is used for all in vitro experiments. (B) Hla lysis of mouse PMNs assessed using LDH assay. Percent cell lysis in untreated control – Young 12.0, aged 11.1 (n=6-8 mice). (C) IL-1β release by LPS-primed human PMNs, after Hla stimulation (n=4-5 humans). (D) IL-1β release by LPS-primed murine PMNs, after Hla stimulation (n=5-6 mice). (E) Skin lesion sizes in young and aged mice 3 days post-Hla (1000 U) challenge (n=7-11 mice). (F) Skin lesion sizes in young or aged mice 3 days post WT or Δ*hla S. aureus* challenge (n=9-10 young mice, 3-4 aged mice). (G-I) Bacterial burden in the skin, spleen, and kidneys of young and aged mice, 3 days post WT or Δ*hla S. aureus* challenge (n=5 young mice, 4 aged mice). Each data point represents an individual mouse. The data are presented as mean ± SEM of biological replicates (A-D). Line represents median (E to I). Two-tailed non-parametric Mann-Whitney U test (A-E), Kruskal-Wallis non-parametric one-way ANOVA test (F-I). Significant P values are shown on the figures; ns, not significant. USA300 WT: *S. aureus* USA300 WT strain SF8300; USA300 Δ*hla*: *S. aureus* USA300 SF8300 Hla isogenic mutant.

### Reduced ADAM10 activity contributes to decreased Hla virulence in senescent mice

Hla mediates various pathologic functions through ligation of ADAM10 on myeloid cells, including neutrophils ^5^. We therefore questioned whether ADAM10 expression or activity differs in neutrophils derived from young and aged mice. Hla transcript and protein levels, as measured by quantitative polymerase chain reaction (qPCR), flow cytometry, and immunoblotting, showed no significant differences between groups (**Figures 2A-D**). Consistent with these findings, Hla binding to neutrophils from young and aged mice was equivalent (**Figures 2E and S2A**). A prior study in Alzheimer’s disease noted that decreased ADAM10 activity contributed to disease pathogenesis ^15^, prompting us to evaluate of ADAM10 enzymatic function in neutrophils and skin tissues. These analyses revealed reduced ADAM10 activity in both neutrophils and skin from aged mice compared to young mice (**Figures 2F-G**), likely reflecting diminished activity in myeloid and skin parenchymal cells, including keratinocytes, a known Hla target ^16^.

**Fig. 2.**
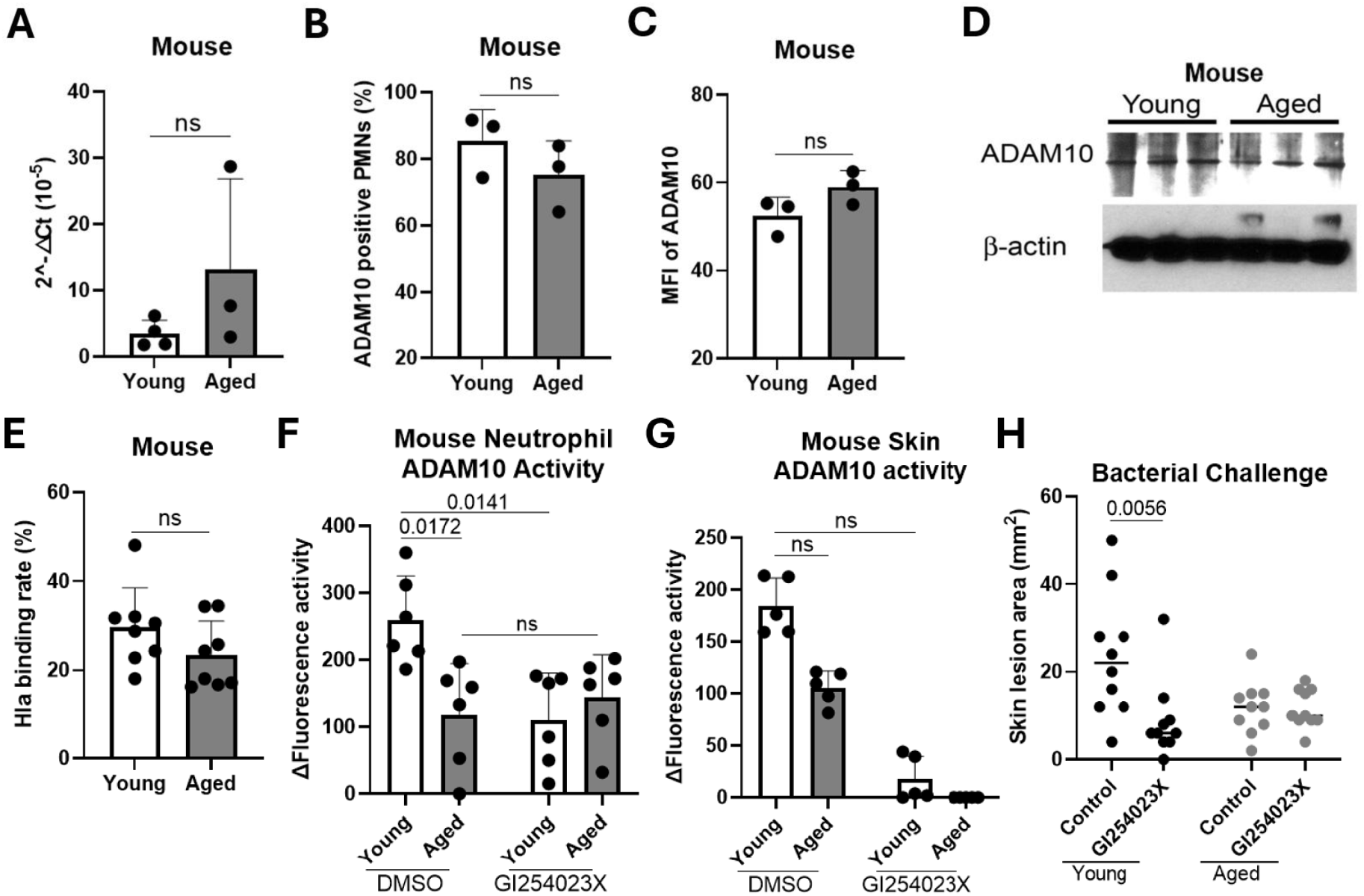
Reduced ADAM10 activity contributes to decreased Hla virulence in aged mice. (A) ADAM10 neutrophil expression derived from young and aged mice, measured by qPCR (n=3-4 mice). (B-C) ADAM10-positive PMNs and mean fluorescence intensity of ADAM10 expression, assessed by flow cytometry (n=3). (D) Neutrophil ADAM10 protein assessed by Western blot (n=3). (E) Binding of Hla to neutrophils assessed by flow cytometry (n=7-8). (F-G) Enzymatic activity of ADAM10 from neutrophil and skin lysates measured with fluorescence-conjugated substrate (n=5-6). (H) Skin lesion sizes in young and aged mice treated with ADAM10 inhibitor GI254023 followed by infection with *S. aureus* (n=10). Each data point represents an individual mouse. The data is presented as mean ± SEM of biological replicates (A-C, E-G). Line represents median (H). Two-tailed non-parametric Mann-Whitney U test (A-C,E), Kruskal-Wallis non-parametric one-way ANOVA test (F-H). Significant P values are shown on the figures; ns, not significant. MFI: mean fluorescence intensity; GI154023X: ADAM10 inhibitor in DMSO; control: DMSO.

ADAM10 contains 4 potential N-glycosylation sites, and glycosylation has been proposed to affect its enzymatic activity ^17^. However, western blot analysis of neutrophils from young and aged mice revealed no detectable difference in N-glycosylation (**Figure S6A**). To investigate the contribution of altered ADAM10 activity to Hla virulence, we treated mice with the ADAM10 inhibitor GI254023 in the context of *S. aureus* subcutaneous infection. Inhibition of ADAM10 significantly reduced lesion sizes in young mice, but had no impact in aged mice, consistent with a functional role for ADAM10 in toxin-mediated pathology only in the young group (**Figures 2H and S3**).

Extending these findings to human samples, we examined neutrophils derived from young and older adult volunteers. As in mice, neutrophils from elderly individuals showed no differences in ADAM10 transcript or protein expression by qPCR, flow cytometry, or western blot (**Figures 3A-D, S4, and S5**) and bound Hla at levels comparable to those from younger donors (**Figures 3E and S2**). However, ADAM10 enzymatic activity was significantly lower in neutrophils from elderly adults, compared to those from young adults (**Figure 3F**). These findings confirm that the age-associated decline in ADAM10 activity is conserved between mice and humans.

**Fig. 3.**
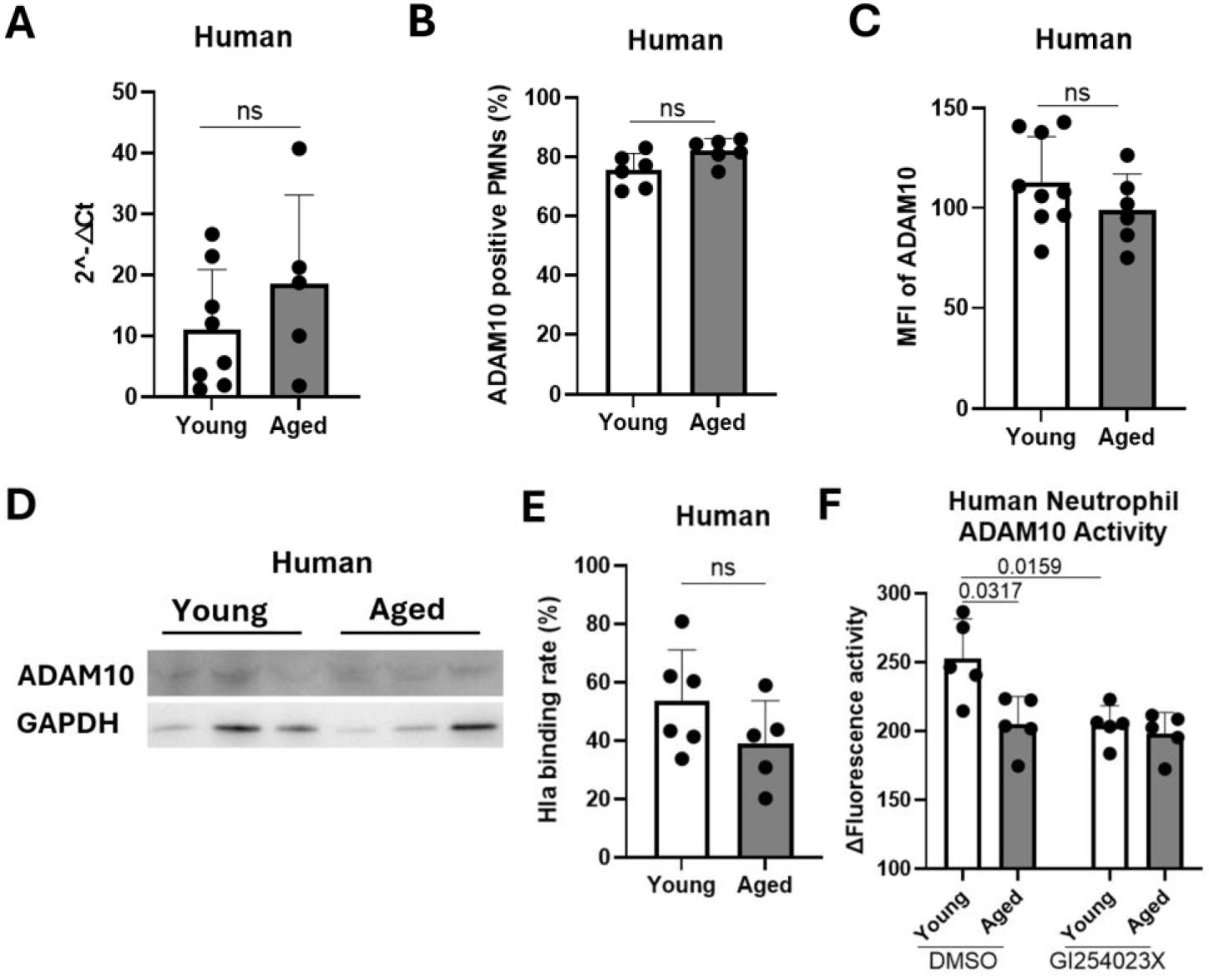
Human PMNs from elderly subjects exhibit reduced ADAM10 activity. (A) ADAM10 expression by neutrophils derived from young and elderly humans, measured by qPCR (n=5-8). (B-C) Percentage and mean fluorescence intensity of ADAM10 expression by neutrophils, assessed by flow cytometry (n =6). (D) Human neutrophil ADAM10 protein assessed by Western blot (n = 3). (E) Binding of Hla to human neutrophils assessed by flow cytometry (n=5-6). (F) Enzymatic activity of ADAM10 from neutrophil lysates, measured with fluorescence-conjugated substrate (n=5). Each data point represents an individual human subject (A-C and E-F). The data is presented as mean ± SEM of biological replicates. Two-tailed non-parametric Mann-Whitney U test (A-C and E), Kruskal-Wallis non-parametric one-way ANOVA test (F). Significant P values are shown on the figures; ns, not significant. MFI: mean fluorescence intensity; GI154023X: ADAM10 inhibitor in DMSO; control: DMSO.

### Hla virulence is attenuated in NSG mice engrafted with bone marrow from aged mice

To assess the relative contribution of myeloid cells to Hla activity in young and aged mice, we performed bone marrow transplantation using donors of different ages (**Figure 4A**). Bone marrow from young or aged mice was transplanted into newborn NOD/SCID/IL2rγ^null^ (NSG) mice, which lack functional innate and adaptive immune systems ^18^. At 16 weeks post-engraftment, recipients of young and aged marrow exhibited equivalent reconstitution of hematopoietic compartments, as evidenced by comparable frequencies of CD45^+^ cells in the spleen and CD11b^+^ myeloid cells in the blood (**Figure 4B-C**). In contrast to cutaneous tissues from naive young and aged mice (**Figure 2G**), ADAM10 activity in whole skin was similar between the two NSG recipient groups (**Figure 4D**), suggesting that ADAM10 function in non-hematopoietic parenchymal tissues (e.g., keratinocytes) was not affected by the age of the donor marrow.

**Fig. 4.**
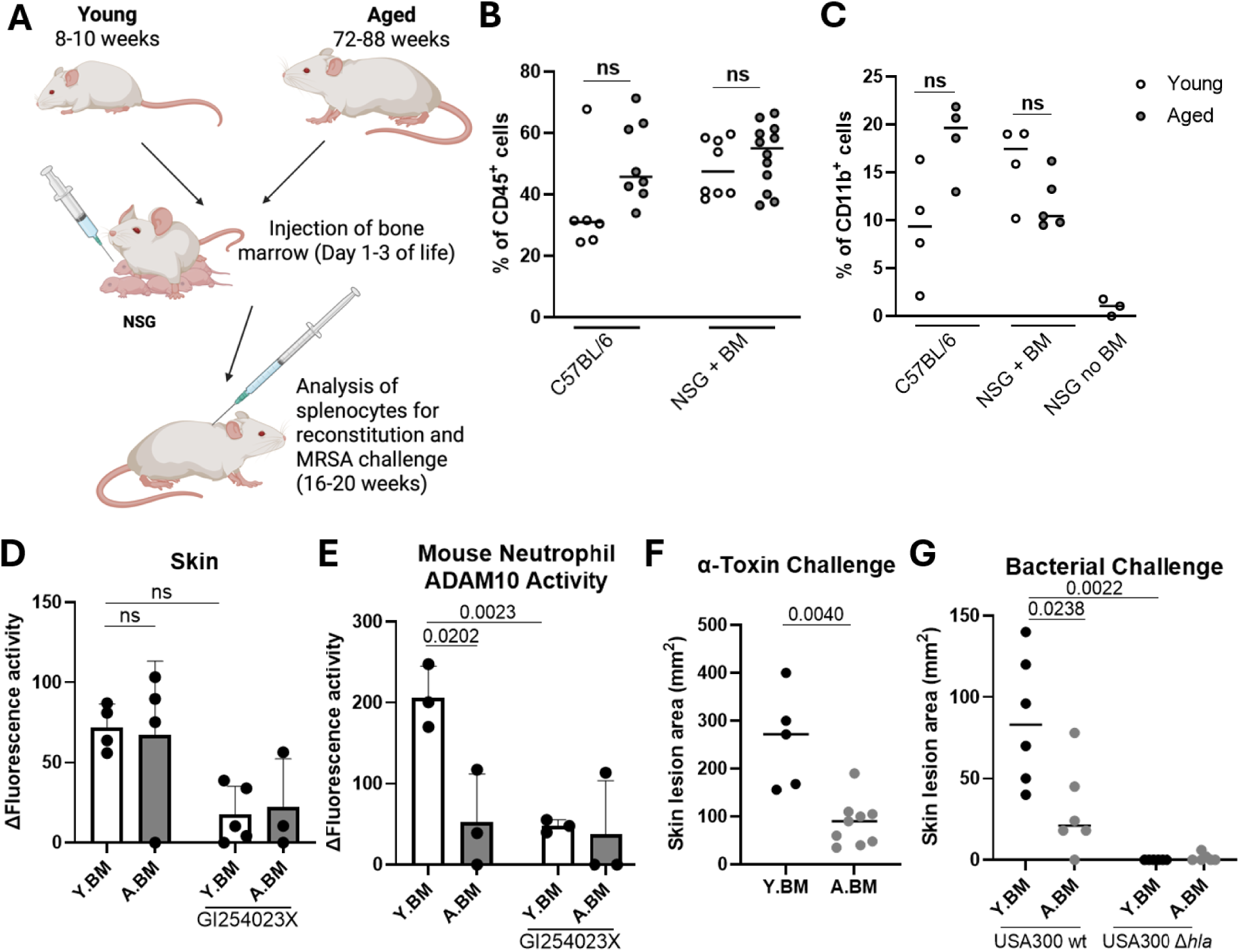
Bone marrow-derived immunocytes contribute to reduced Hla activity in aged mice. (A) Schematic – Bone marrow cells from young or aged mice were injected into 1-to 3-day-old NSG mice. At 16-20 weeks of age, the mice were analysed for reconstitution and then challenged with Hla or *S. aureus*. (B-C) Percentage of splenic CD45^+^ cells and blood CD11b+ cells (n=6-12 for CD45 and n=3-4 for CD11b analysis). (D-E) Enzymatic activities of ADAM10 in neutrophil and skin lysates (n =3-4). (F) Skin lesion sizes 3 days after Hla challenge (n=5-9). (G) Skin lesions sizes 3 days post infection with WT or *Δhla S. aureus* (n=5-6). Each data point represents an individual mouse. Line represents mean (B, C, F, and G). The data is presented as mean ± SEM of biological replicates (D-E). Two-tailed non-parametric Mann-Whitney U test (B, C, F), Kruskal-Wallis non-parametric one-way ANOVA test (D, E, G, H). Significant P values are shown on the figures; ns, not significant. NSG: NOD/SCID/IL2rγ^null^ mouse; Y.BM: NSG recipients of bone marrow from young C57BL/6 mice; A.BM: NSG recipients of bone marrow from aged C57BL/6 mice. GI154023X: ADAM10 inhibitor in DMSO; control: DMSO. The schematic was created in BioRender. Du, X. (2025) https://BioRender.com/h2vi3i1 (A).

However, neutrophils from recipients of aged marrow exhibited significantly reduced ADAM10 activity compared to recipients of young marrow (**Figure 4E), which aligns** with findings in intact aged mice (**Figure 2F**). No difference in ADAM10 glycosylation was observed on neutrophils between the two groups (**Figure S6B**). When challenged with subcutaneous Hla, NSG mice engrafted with aged marrow developed significantly smaller lesions than those receiving young marrow (**Figures 4F, S7A-B**). This finding was further validated in infections with wild-type and Hla-deficient *S. aureus* strains (**Figures 4G, S7C-D**). Together, these findings demonstrate that hematopoietic cells, particularly ADAM10-expressing myeloid cells, are likely important contributors to mediating Hla pathogenesis, and that age-associated reductions in their function are sufficient to attenuate toxin virulence.

### Drug-induced leukopenia limits Hla-mediated pathology

In addition to senescence, we hypothesized that depletion or dysfunction of myeloid cells, such as that induced by chemotherapeutic treatment, would similarly diminish Hla virulence. To test this, we employed a well-characterized murine model of leukopenia using serial moderate-dose cyclophosphamide injections (**Figure 5A**). In this model, wild-type mice infected subcutaneously with either WT or Hla-deficient *S. aureus* developed similar lesion sizes and skin bacterial burdens (**Figures 5B-C**), in contrast to the Hla-dependent pathology observed in immunocompetent controls. In deep tissue sites, cyclophosphamide-treated mice exhibited a trend for increased bacterial burdens that were not dependent on Hla (**Figures 5D-E**).

**Fig. 5.**
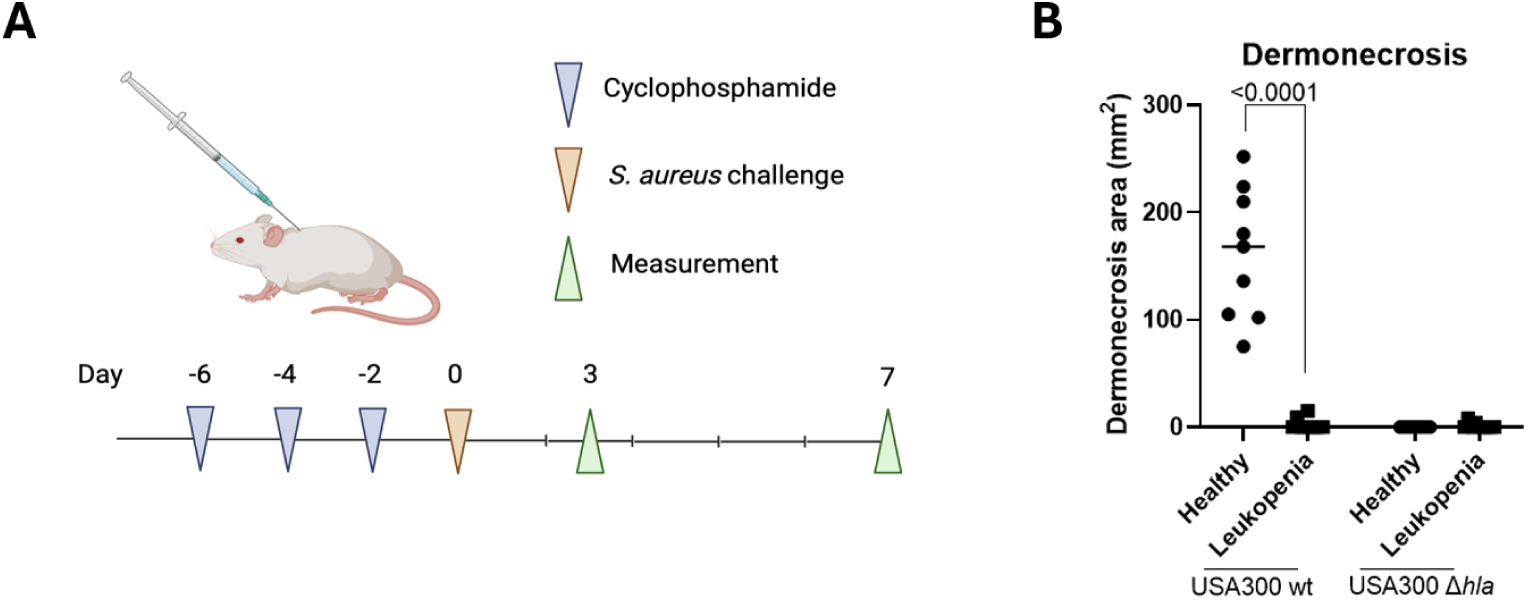
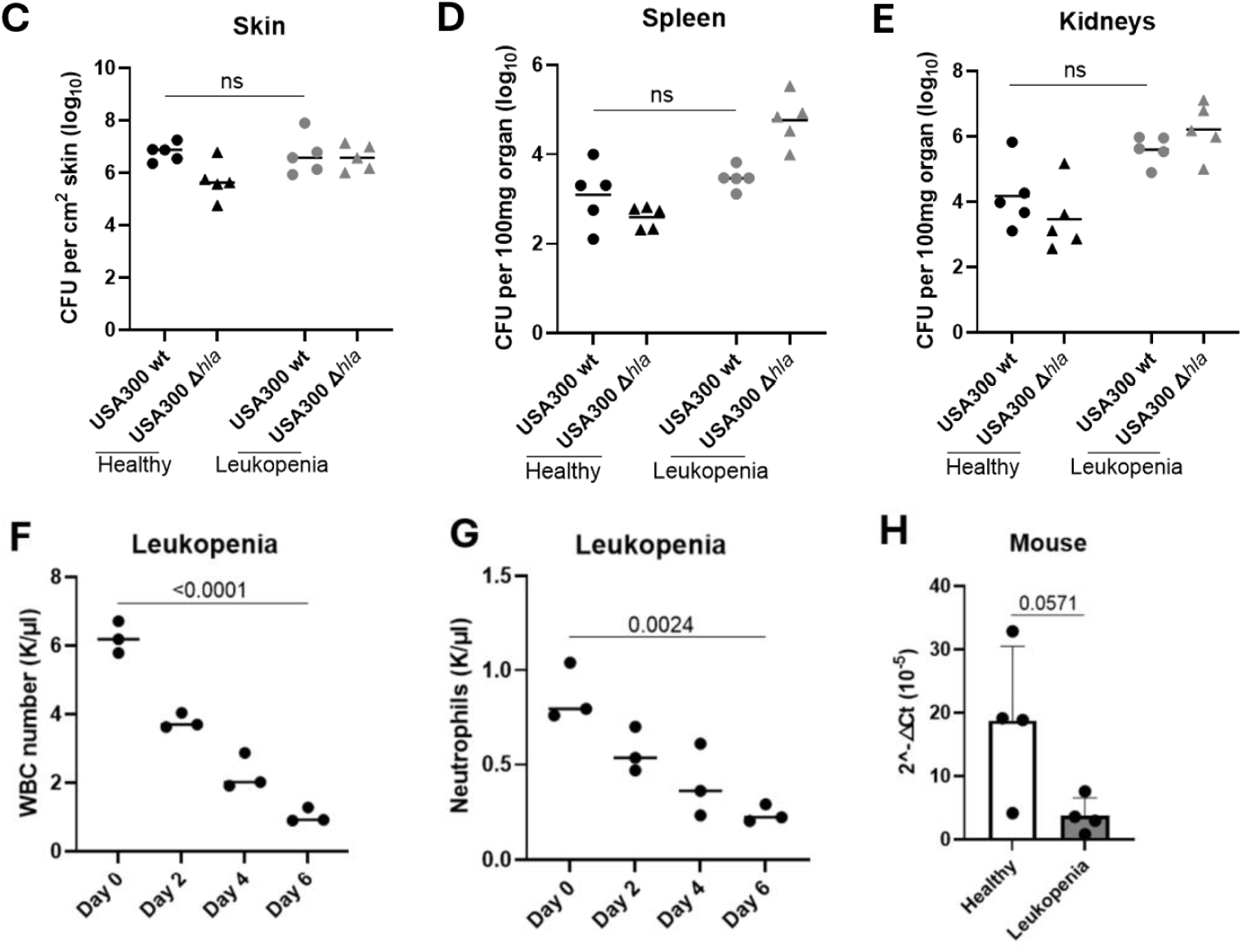
Reduced Hla virulence in mice moderately immunosuppressed with cyclophosphamide. (A) Schematic – Mice were administered moderate doses of cyclophosphamide (75 mg/kg u), then challenged with WT or *Δhla S. aureus*. (B) Skin lesion sizes 3 days post *S. aureus* challenge (n=9). (C) Skin bacterial burden 3 days post *S. aureus* challenge (n=5). (D-E) Spleen and kidney bacterial burden 3 days post *S. aureus* challenge (n=5). (F-G) White blood cell and neutrophil cell count post-cyclophosphamide treatment (n = 3 mice). (H) Neutrophil ADAM10 expression quantified by qPCR (n = 4 mice). Each data point represents an individual mouse (B-H). The data in H is presented as mean ± SEM of biological replicates. Line represents mean (B-G). Two-tailed non-parametric Mann-Whitney U test (H), Kruskal-Wallis non-parametric one-way ANOVA test (B-G). Significant P values are shown on the figures; ns, not significant. The schematic was created in BioRender. Du, X. (2025) https://BioRender.com/pxqt2wr (A).

Flow cytometric analysis confirmed that cyclophosphamide treatment resulted in sustained reductions in circulating white blood cells and neutrophils across multiple time points (**Figures 5F-G**). Unlike aged mice, neutrophils from drug-treated animals did not exhibit a significant change in ADAM10 mRNA, as measured by qPCR (**Figures 5H**). These findings corroborate the reduced pathogenic function of Hla in another critical model of human immunodeficiency, where alternative immune mediators of Hla activity are depleted.

### Active and passive immunizations targeting Hla have modest efficacy in aged mice

Next, we queried whether reduced Hla virulence in aged mice would translate to impaired vaccine efficacy. Using both active immunization with an alpha-toxoid and passive transfer of protective sera, we evaluated protection in young and aged mice following challenge with purified Hla or *S. aureus* (**Figure 6A**). Consistent with prior studies^6,19^, active immunization significantly reduced lesion sizes following challenge with either Hla (**Figures 6B and S8A**) or *S. aureus* in young mice (**Figures 6C, S8C, S8E, and S8F**). In contrast, aged mice showed reduced or no protection under the same vaccination regimen.

**Fig. 6.**
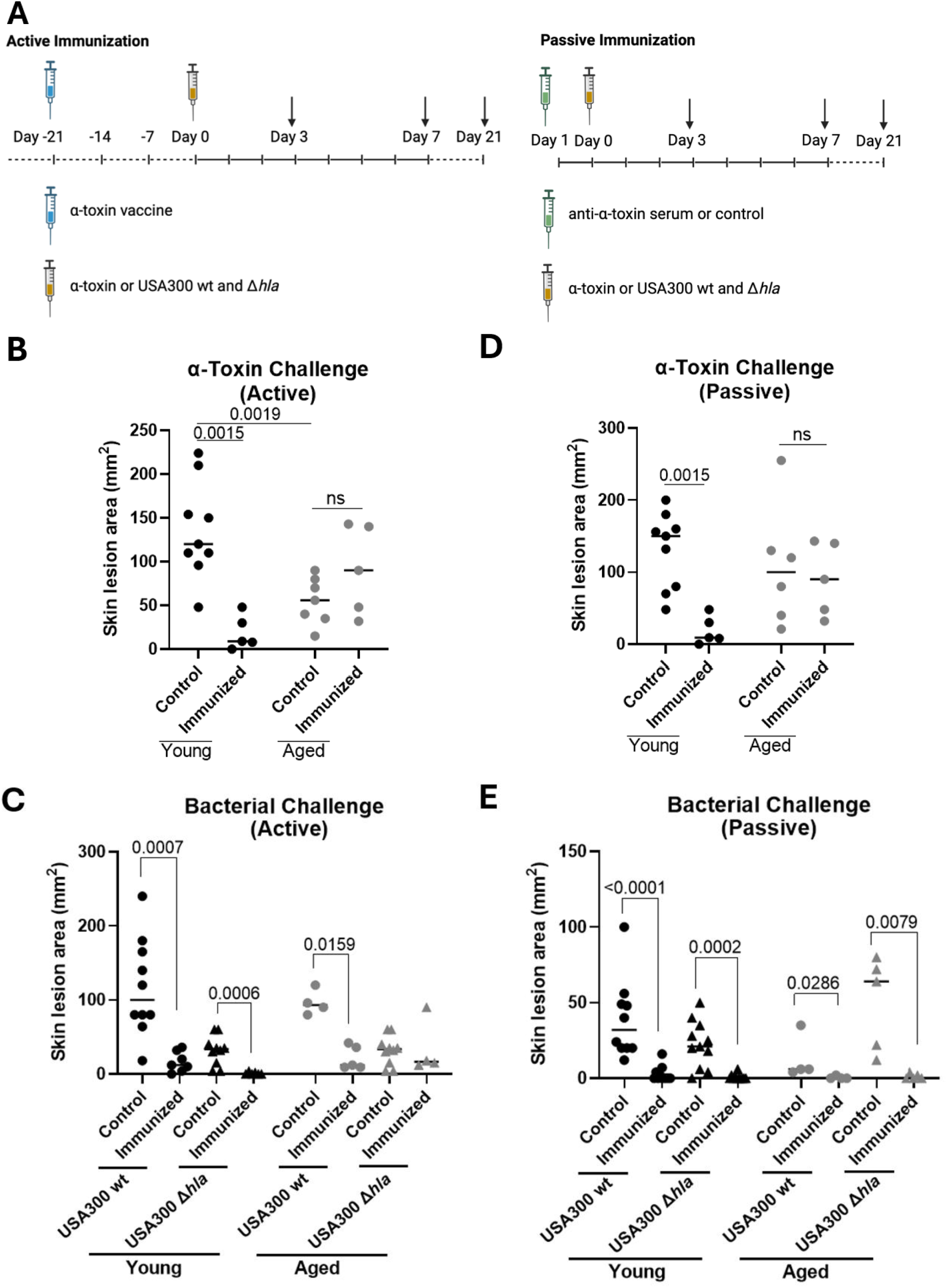
Anti-Hla active and passive immunizations were effective in young but not aged mice. (A) Schematic – Young and aged mice were immunized with alum or alpha toxoid/alum, or administered mouse anti-Hla serum from Hla toxoid injected mice, then challenged with Hla or WT or *Δhla S. aureus*. (B) Active immunization - Skin lesion sizes 7 days after Hla challenge (n=5-9 mice). (C) Active immunization - Skin lesion sizes 7 days after USA300 WT or USA300 Δhla *S. aureus* subcutaneous infection (n=9-11 young, 4-7 aged). (D) Passive immunization - Skin lesion sizes 7 days after Hla challenge (n=5-9). (E) Passive immunization - Skin lesion sizes 7 days after USA300 WT or USA300 Δ*hla S. aureus* subcutaneous infection (n=9-11 young, 4-7 aged). Each data point represents an individual mouse. Line represents mean. Kruskal-Wallis non-parametric one-way ANOVA test (B-E). Significant P values are shown on the figures; ns, not significant. The schematic was created in BioRender. Du, X. (2025) https://BioRender.com/49kavfj, https://BioRender.com/nqcey3h (A).

To further test the impact of host age, we performed passive immunization by adoptively transferring sera from vaccinated young mice into young or aged recipients prior to toxin or *S. aureus* challenge (**Figures 6D-E and S8B, S8D, S8G, and S8H**). Mirroring results from the Suvratoxumab trial, where participants >65 years showed no benefit from anti-Hla antibody therapy ^14^, only young recipients of immune sera were protected, exhibiting significantly smaller lesions after challenge with either Hla or *S. aureus*. Aged recipients showed no protective effect.

Unexpectedly, both active and passive immunization provided some protection against challenge with the Δ*hla S. aureus* challenge in young (active immunization, **Figures 6C and 6E**) and aged (both active and passive immunization, **Figures 6C and 6E**). The mechanism underlying this observation remains unclear, but it did not alter the overall conclusion that Hla-targeted vaccination is less effective in aged hosts. Collectively, these results highlight that the success of anti-toxin vaccination may depend not only on the immunogenicity of the vaccine but also on the preserved pathogenic function of the microbial target in the host.

## Discussion

The current study builds on insights from the Suvratoxumab trial ^14^ to test the hypothesis that the virulence of a major *S. aureus* toxin, Hla, is diminished in aged or immunocompromised hosts, thereby reducing vaccine efficacy. We show that Hla exhibits age-related attenuation in its cytolytic and proinflammatory effects on both murine and human neutrophils. Mechanistically, we find that this reduced pathogenicity is linked to diminished activity, but not expression, of its host receptor, ADAM10. This loss of function impairs the toxin’s ability to damage tissue and incite inflammation. Bone marrow chimera experiments further implicate myeloid cells as key contributors to this shift in Hla activity. In both active and passive immunization models, diminished Hla function translated to loss of vaccine efficacy in aged hosts.

These findings add to existing hypotheses for the repeated failure of *S. aureus* vaccines, including ineffective antigen or adjuvant selection, flawed trial designs, or inappropriate target populations ^13^. Additionally, we previously showed that pre-existing anti-*S. aureus* antibodies could compete with Suvratoxumab for binding, thereby reducing its efficacy in mice ^20^. Here we add another explanation: that diminished virulence of microbial antigen targets in aged or immunocompromised individuals may itself reduce vaccine effectiveness. This concept aligns with the Suvratoxumab trial outcome, where protection was evident in subjects under 65 but not in those older than 65 ^14^.

A second significant point is that microbial virulence is not intrinsic to the pathogen alone, as it reflects an interaction with the host immune system. Direct engagement of host immune receptors and downstream effectors by microbial factors is typically required to induce tissue damage or to evade immune responses. When these host receptors or effectors are altered, due to aging, disease or immunosuppression, virulence factors could lose their pathogenic potential. Consistent with this hypothesis, perturbance of effector functions downstream of the Hla receptor ADAM10, such as neutrophil depletion or impaired neutrophil function caused by moderate-dose cyclophosphamide, was sufficient to reduce Hla-mediated pathology. Based on these findings, we propose that other forms of human immunodeficiency, including diabetes, autoimmune or rheumatologic disease, and therapeutic immunosuppression, may similarly alter the function and clinical impact of microbial virulence factors.

The proposition raises a larger question: is there wider evidence that virulence factor activity is modulated by host immune status? A prior screen by Missiakas and colleagues tested eight *S. aureus* mutants, including Hla and multiple adhesins, in a model of drug-induced severe leukopenia ^21^. In this systemic infection model, seven of eight mutants showed no attenuation in virulence ^21^, suggesting that their pathogenic function was dispensable in immunocompromised hosts. Similarly, our prior work demonstrated that *S. aureus* nuclease lost immune evasion function in aged mice because of reduced neutrophil extracellular trap (NET) formation ^22^. While neither of these studies raised the broader hypothesis across immunocompromised hosts, together they support the hypothesis that virulence factor functions would be frequently altered in immunocompromised settings. These findings challenge current vaccine development strategies, which often rely on virulence factors defined in young immunocompetent animals and do not account for the immune status of target human populations.

Our study has several limitations. Most experiments were conducted in animal models, and although we confirmed reduced Hla activity and diminished Hla responsiveness in neutrophils from aged human donors, the in vivo relevance to human disease remains to be fully validated. We did not evaluate whether microbial gene expression is altered in aged or immunocompromised hosts, though consistent findings across both purified toxin and live bacterial challenge suggest that altered toxin function has a principal role in the observed phenotypes. An unexpected observation was that Hla immunization conferred some protection even against *Δhla S. aureus* challenge, in both young (active immunization) and aged (active and passive immunization) mice. We did not pursue additional experiments to explain the finding, which does not impact our overall conclusion that Hla-targeted vaccination is less effective in aged hosts.

Although diminished Hla function likely contributes to vaccine failure, other factors may play a role, including impaired adaptive immune responses or reduced synergy between transferred antibodies and innate immune effectors in aged hosts ^23^. Although we established the importance of ADAM10 activity and myeloid cells in mediating changes in Hla virulence, we did not dissect all downstream effector pathways. Hla is known to induce NALP3 inflammasome activation and mature IL-1β secretion ^24^, and ADAM10 acts as a sheddase that releases various cytokines to promote inflammation ^25^. These downstream immune responses may be blunted or altered in aged hosts and warrant further study, though they do not affect the primary conclusion of this work. We also acknowledge that age-associated myeloid dysfunctions beyond ADAM10 could contribute to attenuating Hla-mediated damage. For example, aging has been linked to reduced NETosis and impaired neutrophil chemotaxis, which could independently limit dermonecrosis and inflammation ^22,26,27^.

In summary, our findings reveal an apparent paradox: diminished virulence of a major staphylococcal toxin in hosts who are more vulnerable to bacterial infection overall.

Immunocompromised patients frequently receive antimicrobial prophylaxis to reduce the risk of severe infection, and vaccination remains a prevention strategy in those who can mount an effective immune response ^28^. In elderly adults, strategies such as higher antigen dose, improved adjuvants, or alternative delivery routes have been employed or proposed to improve vaccine efficacy ^29^. However, our results caution against reliance on virulence mechanisms defined solely in young, immunocompetent models. In immunocompromised hosts, microbial factors may have reduced, unchanged, or even increased virulence. A deeper understanding of how immune context modulates pathogen behavior is essential to guide rational vaccine design for vulnerable human populations.

## Methods

### Ethics statement

Mouse studies were reviewed and approved by the Institutional Animal Care and Use Committee. Mouse experiments were conducted in accordance with regulations and recommendations on animal experiments cited by the Animal Care Programs at the University of California, San Diego, and Cedars-Sinai Medical Center. Experimentations using human blood were approved by the UCSD Human Research Protection Program. Prior informed consents were obtained from the human subjects. Experimental protocols were approved by the UCSD Biosafety Committee.

### Mice

C57BL/6 young mice (6–12 weeks old) were purchased from Jackson Laboratories. C57BL/6 aged mice (16–22 months old) were either purchased directly from Jackson or purchased at 6 months of age and aged in-house. NOD-SCID IL2rg^−/−^ (NSG) mice were purchased from Jackson Laboratories and used for bone marrow reconstitution. Mice were housed under specific pathogen-free conditions. All animal protocols were approved by the Institutional Animal Care and Use Committee and conducted in accordance with institutional and NIH guidelines. All experiments included both male and female mice unless otherwise indicated.

### Human subjects

Human neutrophils were isolated from freshly drawn peripheral blood of anonymized healthy adult donors from the University of California, San Diego and the San Diego Blood Bank. Young donors were aged 18–35 years; elderly donors were aged 65–85 years. All participants provided informed consent, and the study was approved by the Institutional Review Board of the University of California, San Diego. Donors were screened to exclude immunodeficiencies, ongoing infections, or medication use that could affect immune function.

### Bacterial strain and preparation

Staphylococcal Hla wild-type strain SF8300 (USA300 lineage) and its isogenic knockout SF8300 *Δhla* were generous gifts from Dr. Binh Diep (University of California San Francisco). To preserve phenotypic integrity, frozen glycerol stocks stored at −80°C were freshly streaked onto tryptic soy agar (TSA) or sheep blood agar plates (Hardy Diagnostics) and incubated at 37°C for 16-18 hours. Hemolytic zones were assessed to confirm Hla expression. For mouse infections and in vitro assays, bacteria were cultured in tryptic soy broth (TSB, BD Difco) at 37°C with agitation at 200 rpm. The bacteria were grown to mid-logarithmic phase (OD600 ≈ 0.7–1.0), harvested by centrifugation at 4,000×g for 10 minutes at 4°C, and washed twice with sterile Dulbecco’s phosphate-buffered saline (DPBS; Gibco). Final bacterial suspensions were adjusted to the desired CFU concentration using optical density (OD) calibration and confirmed by serial dilution and plating on TSA.

### Hla preparation and inactivation

Purified *S. aureus* alpha-toxin (Hla, Sigma-Aldrich) was reconstituted in sterile DPBS at a stock concentration of 10,000 units/ml and aliquoted for single use to minimize freeze-thaw degradation. For passive vaccine preparation, Hla toxoid was generated by incubating Hla with 0.4% methanol-free formaldehyde (Thermo Fisher) at 37°C for 48 hours with gentle agitation. Complete inactivation was confirmed by loss of hemolytic activity on sheep blood agar and cytotoxicity assays in vitro. Endotoxin levels were confirmed to be less than 0.1 EU/μg protein using a Limulus Amebocyte Lysate assay (Lonza).

### Active and passive immunization

For active immunization, mice were anesthetized briefly with isoflurane and injected subcutaneously with 1000 U of Hla toxoid in 100 μl sterile DPBS into the left flank of mice using a 30G insulin syringe on day 21 before challenge with either 1000 U of native Hla or live *S. aureus* (USA300 WT or *Δhla* mutant). Measurement of skin lesions and bacterial burden was conducted on day 3 and day 7 post-challenge.

For passive immunization, Hla toxoid was emulsified 1:1 (v/v) with AddaVax (InvivoGen) before injection. Booster doses were given on day 14 and day 28. Serum was harvested on day 35 pooled, tested for Hla neutralizing activity, and stored at −80°C for passive immunization. A total of 1 mL of 1:2 diluted immune or control serum was administered intraperitoneally 24 hours before challenge with either native Hla (1000 U/100 μl DPBS) or live *S. aureus* (USA300 WT or Δ*hla* mutant, 2-3×10^8^ CFU/ml in 100 μl DPBS). Measurement of skin lesions and bacterial burden was conducted on day 3, 7, and 21 post-challenge.

### Subcutaneous skin infection model

Subcutaneous infection was performed following an established protocol ^30^. Mice were anesthetized with isoflurane and inoculated subcutaneously on the back with either 100 μL of Hla (1000 U) or 2–3× 10^8^ CFU/mL *S. aureus* bacterial suspension in DPBS. Animals were monitored daily and skin lesions were measured using a caliper. On day 3, tissue samples (skin, kidney, spleen) were harvested aseptically, and homogenized in 0.5 ml sterile DPBS. Tissue homogenates were serially diluted, plated on TSA agar plates, and incubated overnight (16-18 hours) for CFU enumeration.

### Bone marrow transplantation

Donor bone marrow cells were collected from femurs and tibiae of 2-month-old (young) or 18-22-month-old (aged) C57BL/6 mice. Cells were flushed with sterile DPBS containing 2% FBS and filtered through a 70 μm strainer. Red blood cells were lysed with lysis buffer, and nucleated cells were counted using trypan blue exclusion. A total of 1× 10^5^ viable bone marrow cells were injected intrahepatically into 1–3-day-old NSG pups (Jackson Labs) using a 30G needle under sterile conditions. Mice were weaned at 3 weeks of age and monitored for 16 weeks.

Reconstitution of myeloid lineages was confirmed by flow cytometry for CD45^+^ and CD11b^+^ cells in spleen and peripheral blood.

### Cyclophosphamide-induced leukopenia of mice

Cyclophosphamide monohydrate (Sigma) was dissolved in sterile DPBS. Mice were administered three doses of cyclophosphamide (75 mg/kg body weight) every 48 hours via intraperitoneal injection. DPBS without cyclophosphamide monohydrate was used as a negative control. One day after the third drug injection, the mice were subcutaneously challenged with *S. aureus*.

### Neutrophil Isolation

Murine bone marrow neutrophils were isolated using MojoSort Mouse Neutrophil Isolation Kit (BioLegend) following the manufacturer’s protocol. Human neutrophils were isolated from 30 mL whole blood samples from volunteers at the University of California, San Diego and San Diego Blood Bank. Human neutrophils were isolated and purified using one-step PolymorphsTM (Fresenius Kabi Norge AS) following the manufacturer’s instructions. Cells were resuspended in RPMI 1640 (Gibco) containing 10% FBS.

### Cytotoxicity assay

Neutrophils were plated in 96-well flat-bottom tissue culture plates (Corning) at 5×10^6^ cells per well in 100 μl Hanks’ Balanced Salt Solution (HBSS without Ca^2+^ and Mg^2+^, Invitrogen) with 0.5% BSA. Cells were incubated with 10 U/ml native Hla or vehicle control at 37°C with 5% CO_2_ for 3 hours. Cell lysis was quantified by measuring lactate dehydrogenase (LDH) release using a colorimetric LDH Cytotoxicity Assay Kit (Takara Clontech) according to the manufacturer’s instructions. Absorbance was read at 490 nm using a microplate reader SMARTREADER 96 (ACURIES). Cytotoxicity was calculated as the percentage of maximum lysis.

### Determination of IL-1β in neutrophil culture supernatants

Enzyme-Linked Immunosorbent Assay (ELISA) was performed using an IL-1β ELISA MAX Deluxe Set (BioLegend) using supernatants from Hla-treated mouse and human neutrophils. Neutrophils purified from mouse bone marrow or human blood samples were incubated with LPS (200 ng/ml) (eBioscience) for 4 hours, then treated with Hla (10 U/ml) for 2 hours at 37°C. The supernatants were then tested for IL-1β levels following the manufacturer’s instructions.

Absorbance was measured at 450 nm using a microplate reader SMARTREADER 96 (ACURIES). Cytokine concentrations were determined by interpolation from a standard curve generated with known concentrations of recombinant IL-1β.

### Quantitative real-time PCR

Total RNA was isolated from tissue or neutrophil lysates using the RNeasy Mini Kit (Qiagen), treated with TURBO DNA MiniPrep kit (Invitrogen) for DNA removal, and reverse-transcribed with SuperScript III (Invitrogen). qPCR was performed using SYBR Green Master Mix (Bio-Rad) on a CFX96 Touch Real-Time PCR Detection System (Bio-Rad). Relative mRNA levels were normalized to Mouse 18s or GAPDH and calculated using the 2^-ΔCt method. Primer (Integrated DNA Technologies) sequences of ADAM10 and housekeeping genes are as follows:

For mouse ADAM10

Forward ADAM10: 5’-TCATGGTGAAACGCATAAGAATCA-3’

Reverse ADAM10: 5’-CCAGACCAAGTACGCCATCA-3’

Ms 18s Forward: 5’-GCCGCTAGAGGTGAAATTCTT-3’

Ms 18s Reverse: 5’-CGTCTTCGAACCTCCGACT-3’

For human ADAM10

ADAM10_F Sequence: 5’-CTGGCCAACCTATTTGTGGAA-3’

ADAM10_R Sequence: 5’-GACCTTGACTTGGACTGCACTG-3’

GAPDH_ F Sequence: 5’-GAAGGGCTCATGACCACAGTCCAT-3’

GAPDH_R Sequence: 5’-TCATTGTCGTACCAGGAAATGAGCTT-3’

### ADAM10 activity assay

ADAM10 enzymatic activity was measured using the ADAM10 Fluorogenic Assay Kit (BPS Biosciences). Neutrophil lysates were prepared by lysing cells in RIPA lysis buffer (Millipore Sigma) with cOmplete protease inhibitor cocktail (Sigma) on ice for 20 minutes, followed by a high-speed centrifugation at 14,500×g for 10 minutes at 4°C. Neutrophil lysates or tissue homogenates (5 μg protein per reaction) were used for the reaction in each well of 96-well plates with or without the ADAM10-specific inhibitor GI254023. Fluorescence (Ex/Em: 485/538 nm) was recorded after a 5-minute reaction at room temperature using a fluorescence microplate reader.

### Flow cytometric analysis of neutrophil ADAM10 surface expression

Neutrophils, with or without Hla pretreatment, were washed with FACS buffer (DPBS + 1% BSA), blocked with anti-CD16/CD32 Fc block (BioLegend), and then stained with fluorophore-conjugated monoclonal antibodies against CD11b, Ly6G or CD15, and ADAM10 (BioLegend), to detect surface ADAM10 on neutrophils. UltraComp eBeads Compensation beads (Invitrogen) were used for single fluorescence control. Flow cytometry was performed on a BD FACSCanto II flow cytometer and analyzed using FlowJo_v10.9.0_CL.

### Hla binding assay

Hla binding to neutrophils was performed using Hla toxoid labeled with Alexa Fluor 647, using Alexa Fluor 647 protein labeling kit (Invitrogen). After co-incubation with the fluorophore-labeled Hla toxoid at 37°C for 15 minutes, the cells were washed twice and stained with fluorophore-conjugated monoclonal antibodies (BioLegen) against CD11b, Ly6G, or CD15 for recognition of neutrophils. Flow cytometry was performed on a BD FACSCanto II flow cytometer and analyzed using FlowJo_v10.9.0_CL.

### Deglycosylation and western blotting

Western blotting of whole-cell lysates was performed on SDS-PAGE, probed for ADAM10 and β-actin or GAPDH, and visualized with HRP-conjugated secondary antibodies and chemiluminescent substrate. For deglycosylation, lysates were treated with PNGase F (New England BioLabs) for 1 hour at 37°C. For Western blot, the protein samples were boiled with NuPAGE LDS Sample Buffer (Invitrogen), separated by SDS-PAGE with NuPAGE 10% Gel (Invitrogen), and transferred onto PVDF membranes using Power Blotter Select Transfer Stacks (Invitrogen). Membranes were blocked with 5% non-fat skim milk (bioWORLD) in Tris-Buffered Saline with Tween 20 (Invitrogen), probed with antibodies against ADAM10 and β-actin or GAPDH (Santa Cruz), followed by HRP-conjugated goat-anti-rabbit secondary antibodies (Invitrogen). The bands on the membrane were visualized using SuperSignalTM West Femto Maximum Sensitivity Substrate (Thermo Scientific).

### Statistics

The sample size, statistical methods, and relevant details are provided in the figure legends, main text, or STAR Methods section. GraphPad Prism 10.4.1 was used for all statistical analyses. Two-group analyses used unpaired Student’s t test (two-tailed tests) or non-parametric Mann-Whitney U-test. Comparisons of multiple groups used one-way ANOVA, with the Kruskal-Wallis test for non-normality. Data were presented as mean ± standard error of the mean (SEM), unless otherwise indicated.

## Data availability

All data supporting the findings of this study are available within the paper and supplementary information.

## Acknowledgments

We thank Dr. Qiongyu Chen from the Department of Hematology for performing the complete cell count for mouse blood. The study received funding from the National Institute of Health grants R01AI127406 (G.Y.L), R01AI144694 (G.Y.L), and R01AI181321 (G.Y.L). We used ChatGPT to assist with grammar and language refinement of the manuscript.

## Author contributions

X.D. and C.W.T. performed most experiments. X.D., C.W.T., and G.Y.L. designed the study. E.B., H.G., and J.S. participated in collecting the human samples and the *in vitro* experiments. C.M.T., I.A.H., C.G., and B.L. participated in the *in vivo* mouse assays and infection experiments. X.D., C.W.T. and GYL analyzed the data. X.D., C.W.T., V.N. and G.Y.L. wrote the manuscript, with all authors providing significant input.

## Additional information

## Supplementary information

Figures S1 to S8

Correspondence and requests for materials should be addressed to Ching-Wen Tseng or George Y. Liu.

## Supplementary materials

**Fig. S1.**
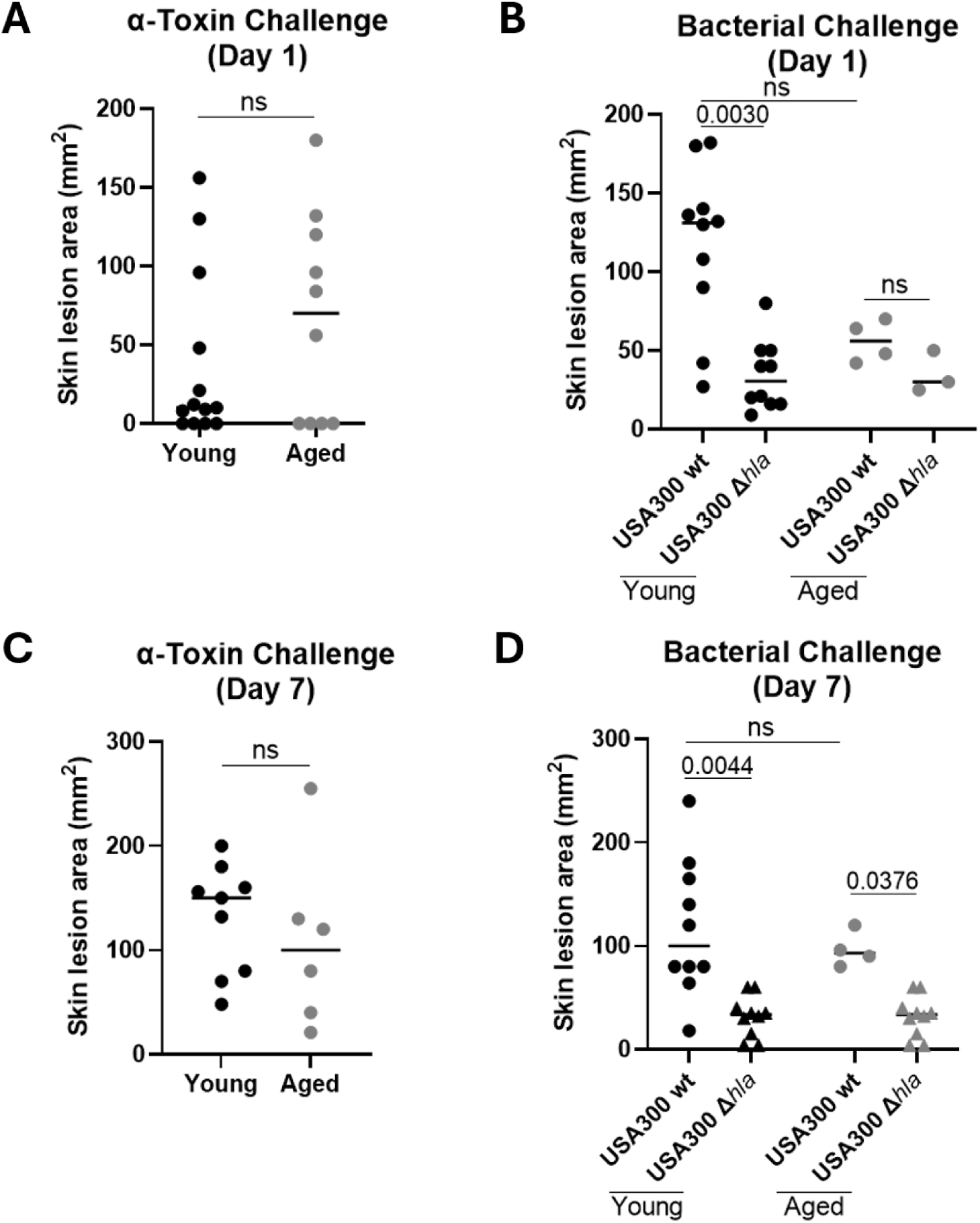
Skin lesion sizes at day 1 and day 7 post-Hla or *S. aureus* challenge of young and aged mice. (A) Hla-induced skin lesions at day 1 post-challenge in young and aged mice. (B) Skin lesions induced at day 1 post-challenge by WT or *Δhla S. aureus* in young and aged mice. (C) Hla-induced skin lesions at day 7 post-challenge in young and aged mice. (D) Skin lesions induced by WT or *Δhla S. aureus* at day 7 post-challenge in young and aged mice. Each data point represents an individual mouse. Line represents mean. Two-tailed non-parametric Mann-Whitney U test (A), Kruskal-Wallis non-parametric one-way ANOVA test (B).

**Fig. S2.**
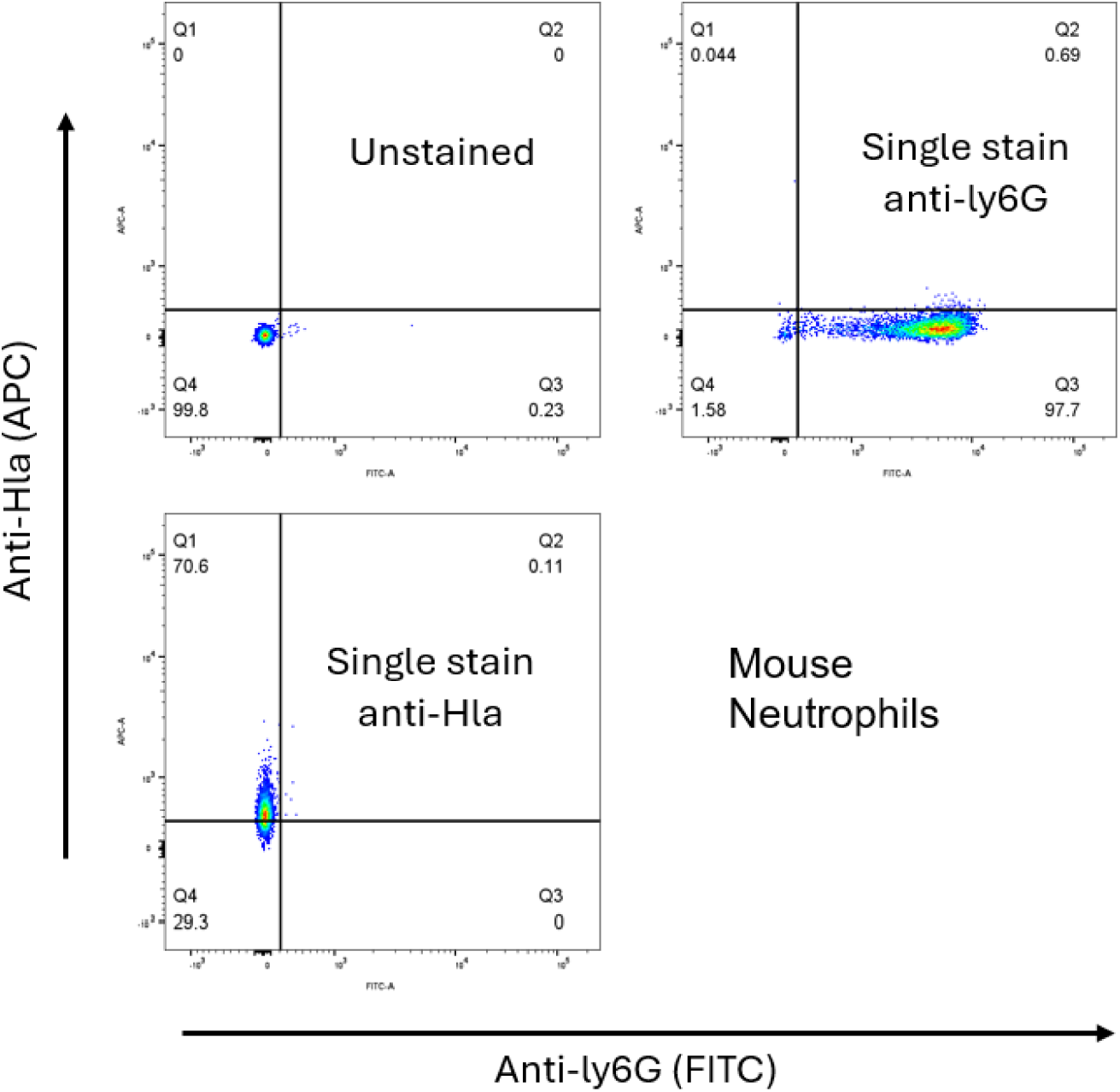
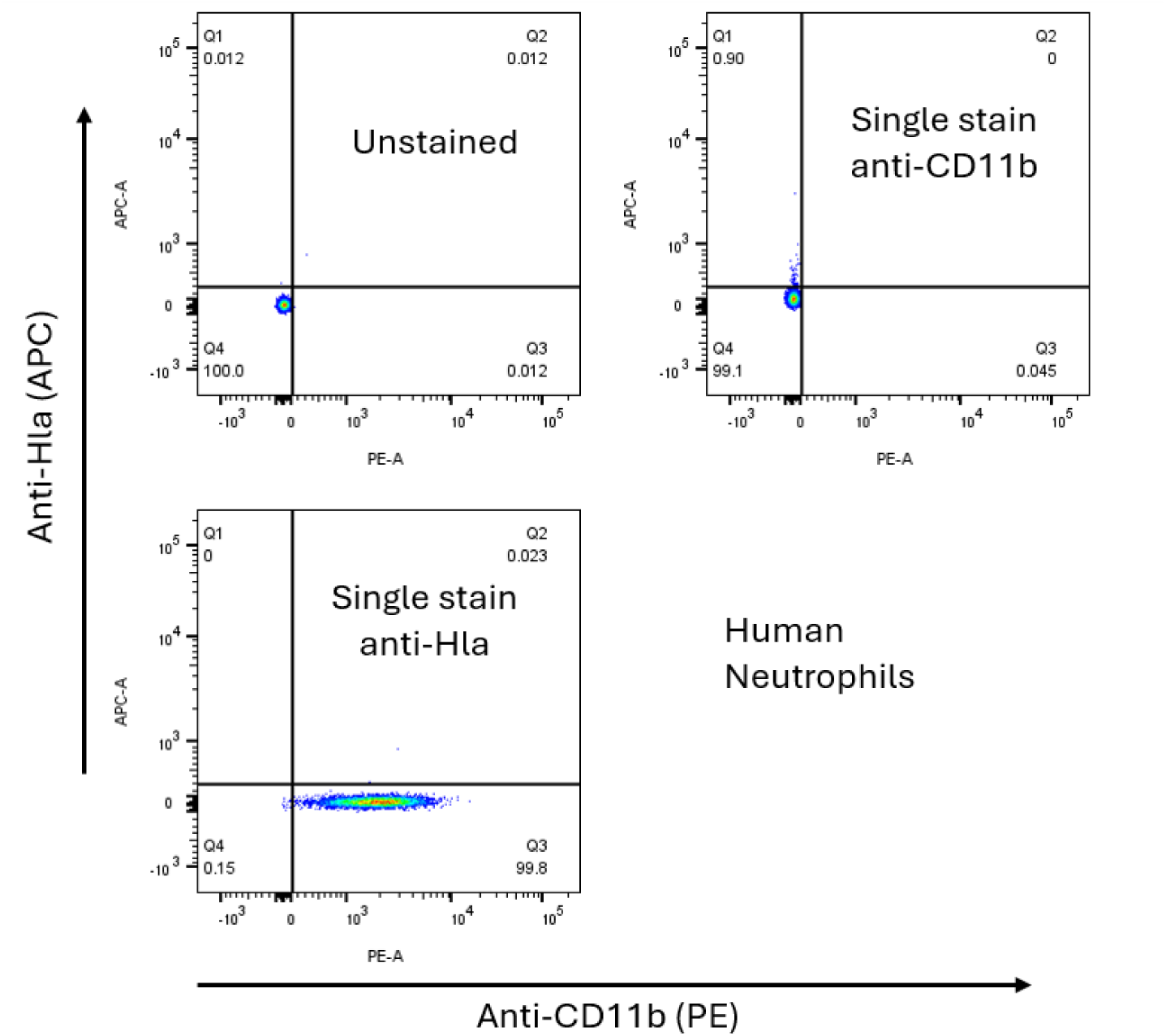
Flow Cytometer Gating Strategy for assessment of ADAM10 binding to PMNs from mice (upper) and humans (lower). Neutrophils purified from young and aged mice or human volunteers were untreated or Hla pretreated for 30 minutes, then stained with both fluorescence-conjugated antibodies against Hla, CD markers for neutrophils.

**Fig. S3.**
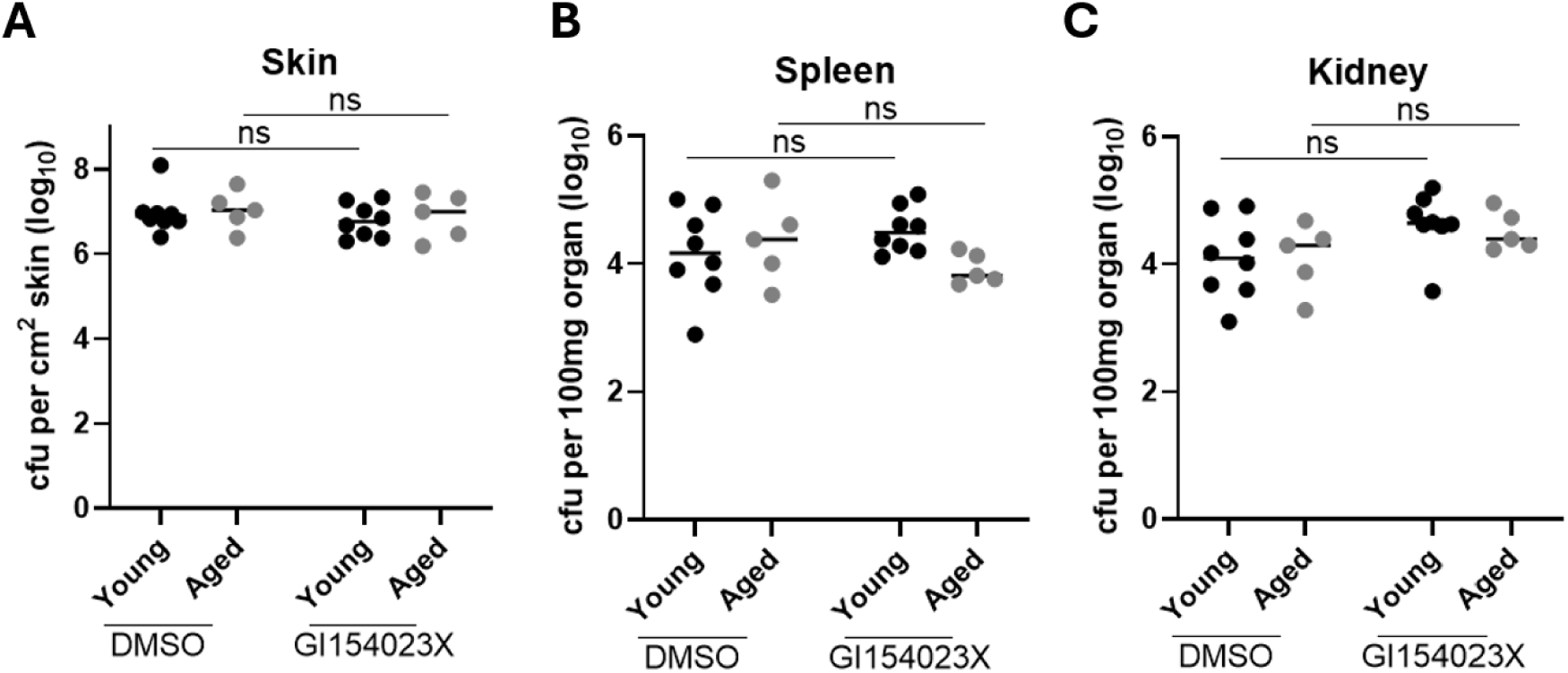
Bacterial burden in young and aged mice treated with ADAM10 inhibitor and challenged with *S. aureus*. (A-C) Young and aged mice were treated with DMSO carrier or ADAM10-specific inhibitor GI154023X, then challenged with WT *S. aureus*. Shown are bacterial burden from the skin (A), spleen (B), and Kidneys (C), 3 days post-infection. Line represents mean. Kruskal-Wallis non-parametric one-way ANOVA test (A-C). *S. aureus*: USA300 WT SF8300.

**Fig. S4.**
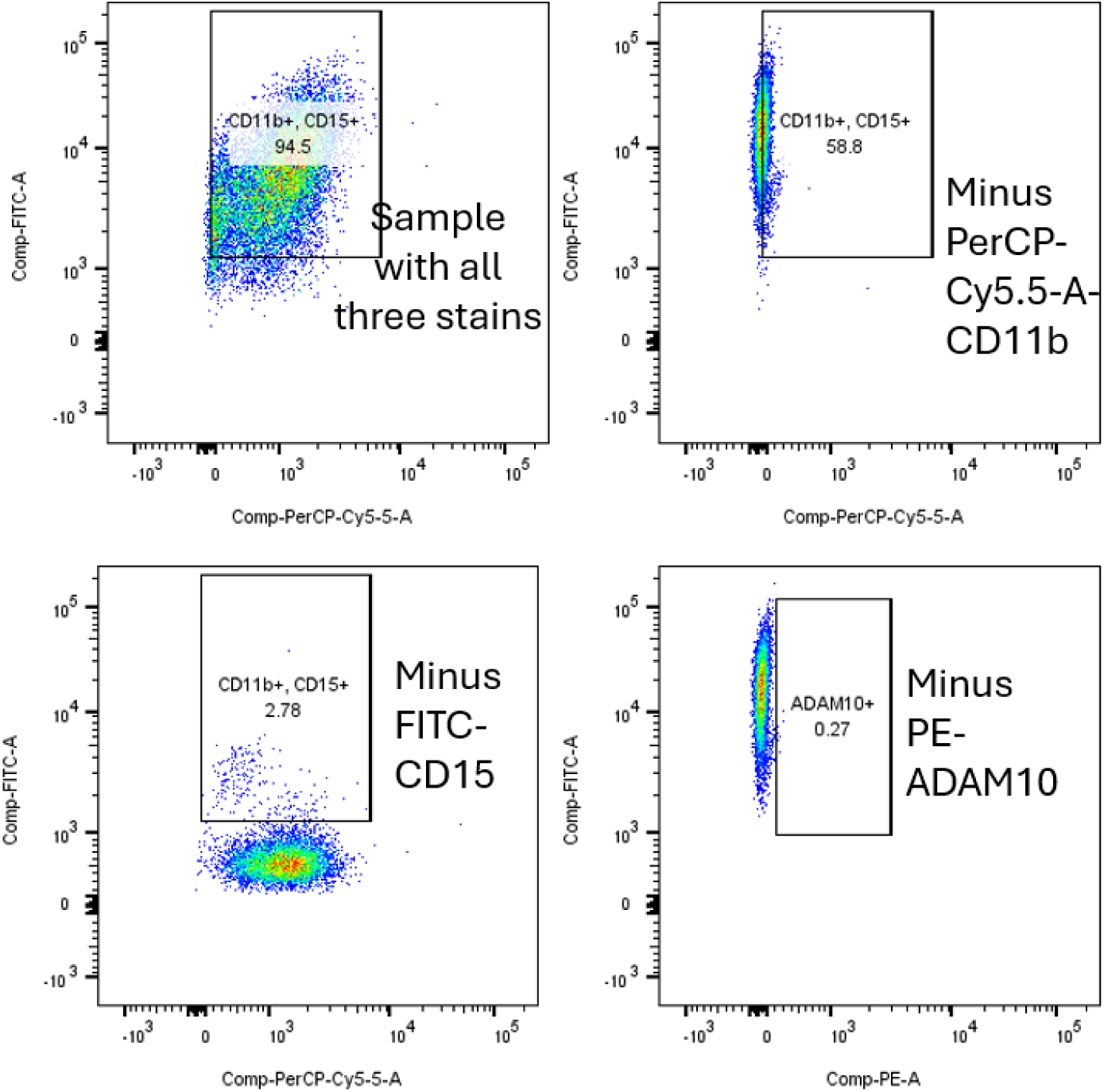
Flow Cytometer gating strategy for assessment of ADAM10 expression by human PMNs. Neutrophils purified from young and aged healthy human volunteers were untreated or Hla pretreated for 30 minutes and stained with both fluorescence-conjugated antibodies against CD markers for neutrophils and ADAM10.

**Fig. S5.**
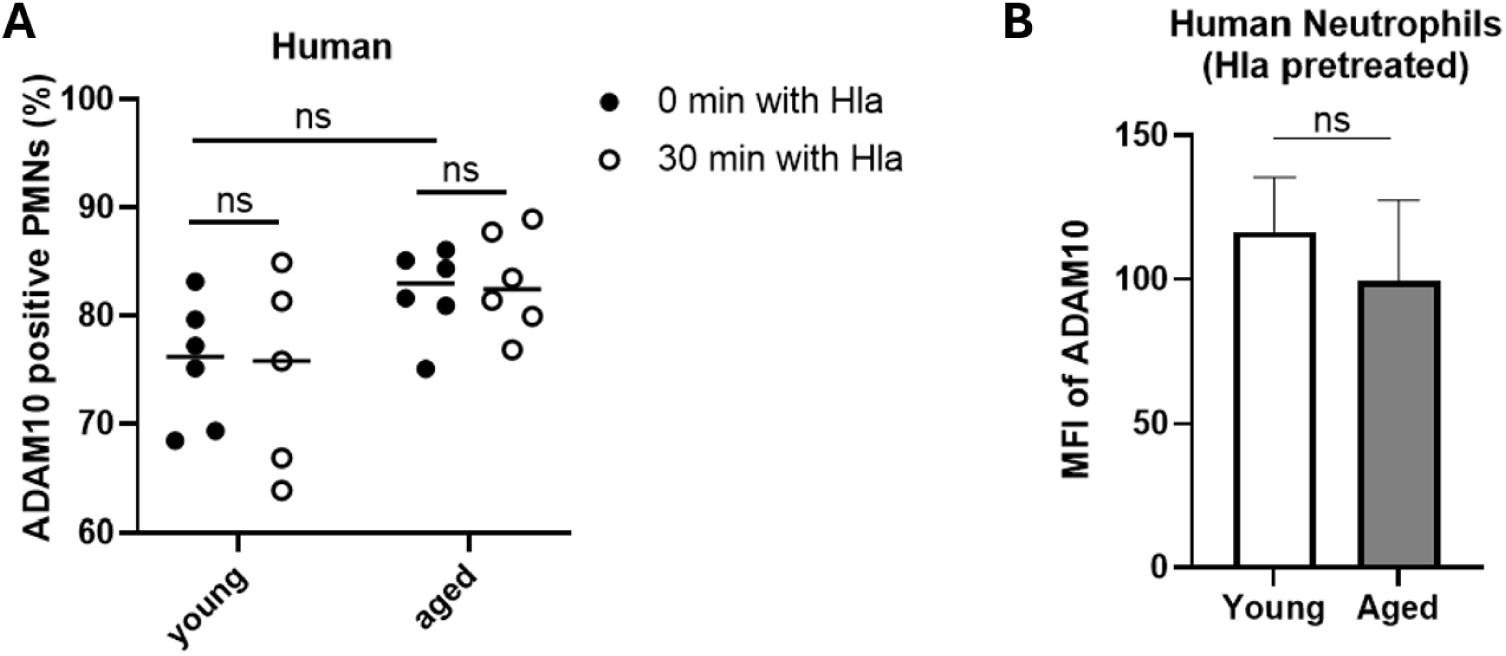
ADAM10 expression by human neutrophils following Hla treatment. (A) Percentage of ADAM10-positive neutrophils 30 minutes after Hla treatment (n=5-6). (B) Mean fluorescence intensity (MFI) of ADAM10 expression 30 minutes after Hla treatment (n=6). Line represents mean (A); the data is presented as mean ± SEM of biological replicates (B). ns, not significant, two-tailed non-parametric Mann-Whitney U test (B); Kruskal-Wallis non-parametric one-way ANOVA test (A).

**Fig. S6.**
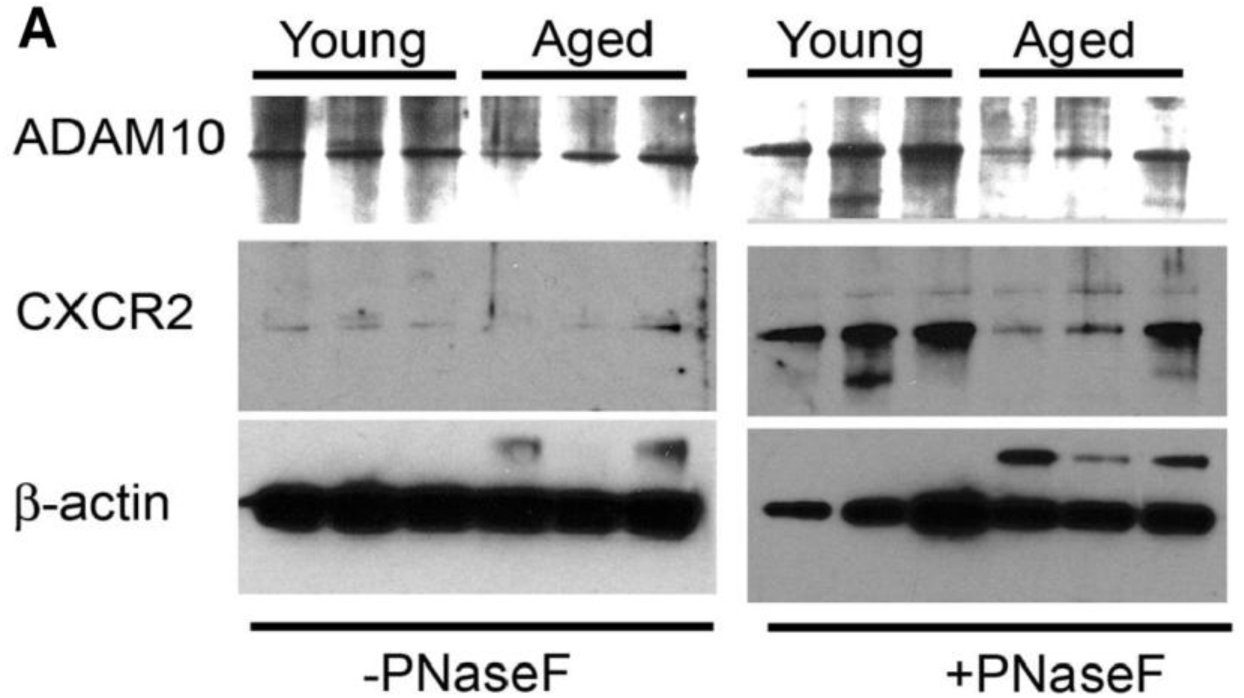
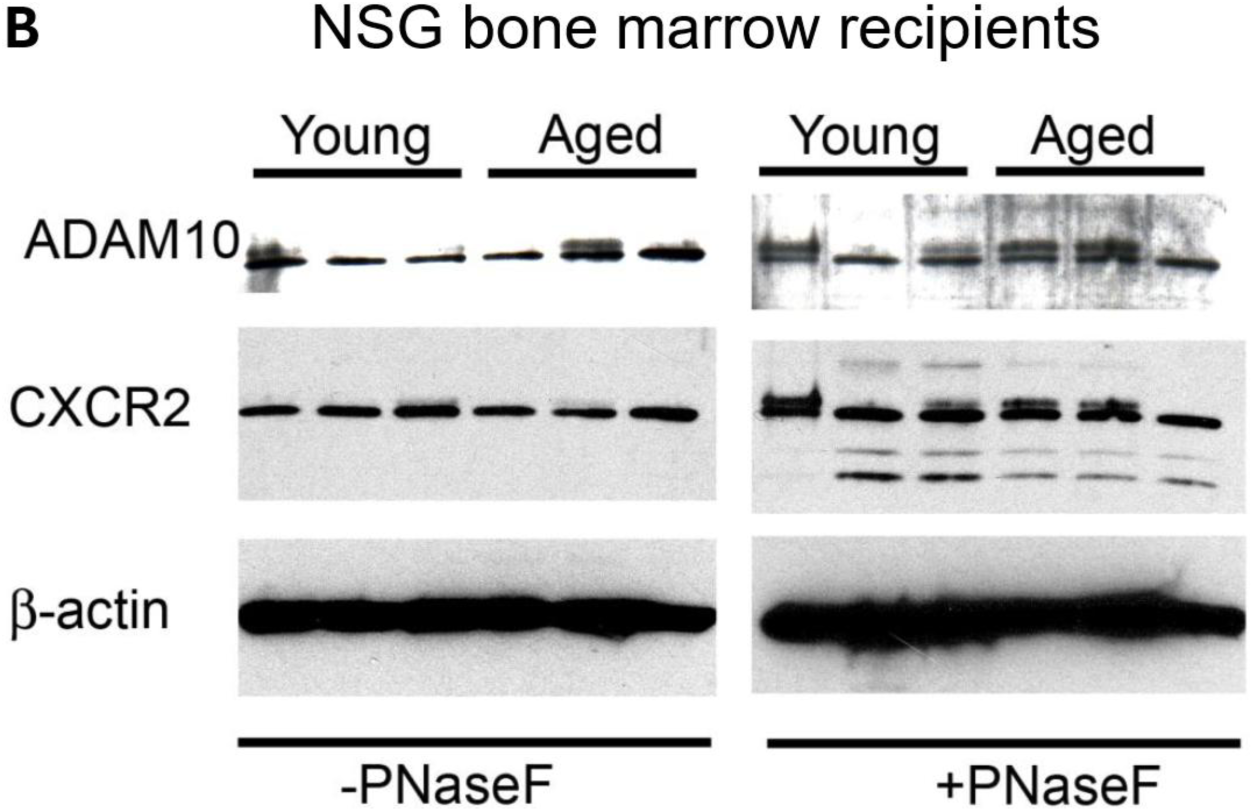
No difference in ADAM10 glycosylation in PMNs isolated from young and aged mice or from NSG mice transplanted with young or old bone marrow. (A-B) PMNS were isolated from young and aged mice (n=3) or NSG mice transplanted with young or old bone marrow (n=3). Shown are the Western blots of ADAM10 protein with or without Peptide-N-Glycosidase F treatment, from young and aged mice (A), and from NSG recipients of young or old bone marrow (B). The β-actin was used as a loading control. CXCR2 was used as a positive control for the N-glycosidase F PNaseF in the treatment.

**Fig. S7.**
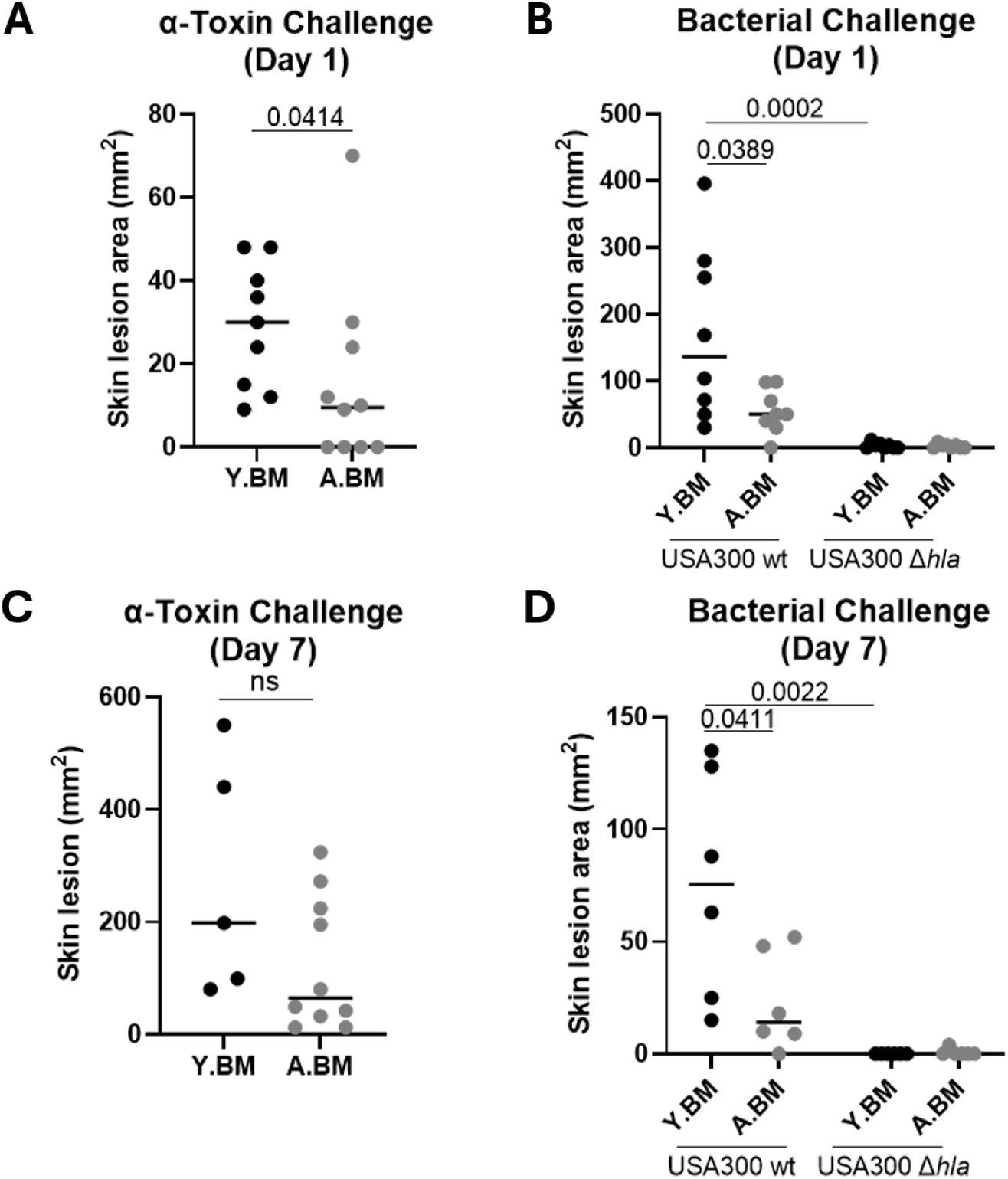
Quantitation of skin lesions after Hla or *S. aureus* challenge in recipients of young and aged bone marrow. NSG mice received bone marrow from young or aged mice. At 16 weeks of age, the mice were challenged with Hla, WT or *Δhla S. aureus*. (A-B) Skin lesion sizes 1 and 7 days after Hla challenge. (C-D) Skin lesion sizes 1 and 7 days after *S. aureus* challenge. Line represents mean. Two-tailed non-parametric Mann-Whitney U test (A-D). Y.BM: NSG recipients of bone marrow from young C57BL/6 mice; A.BM: NSG recipients of bone marrow from aged C57BL/6 mice.

**Fig. S8.**
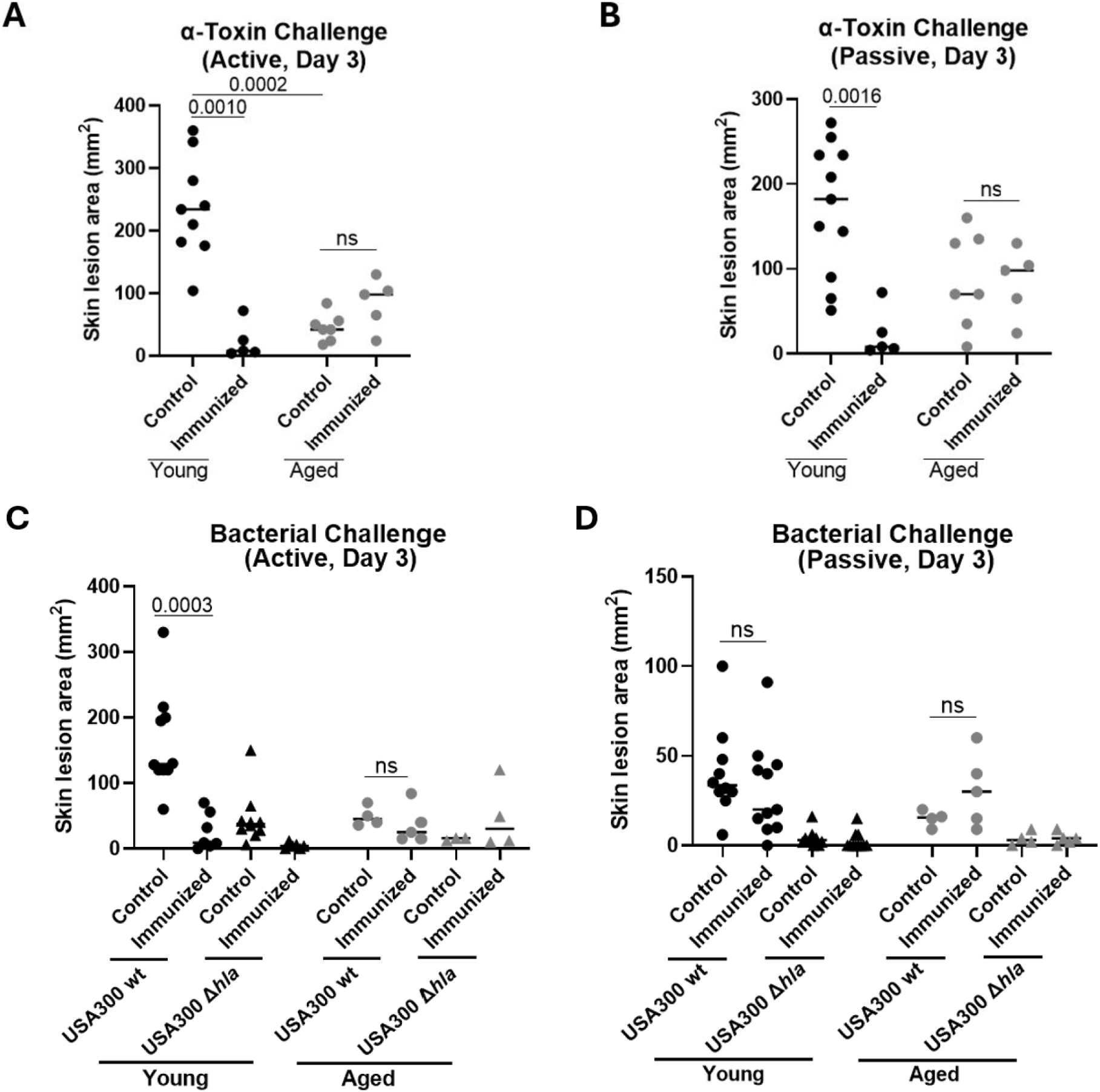
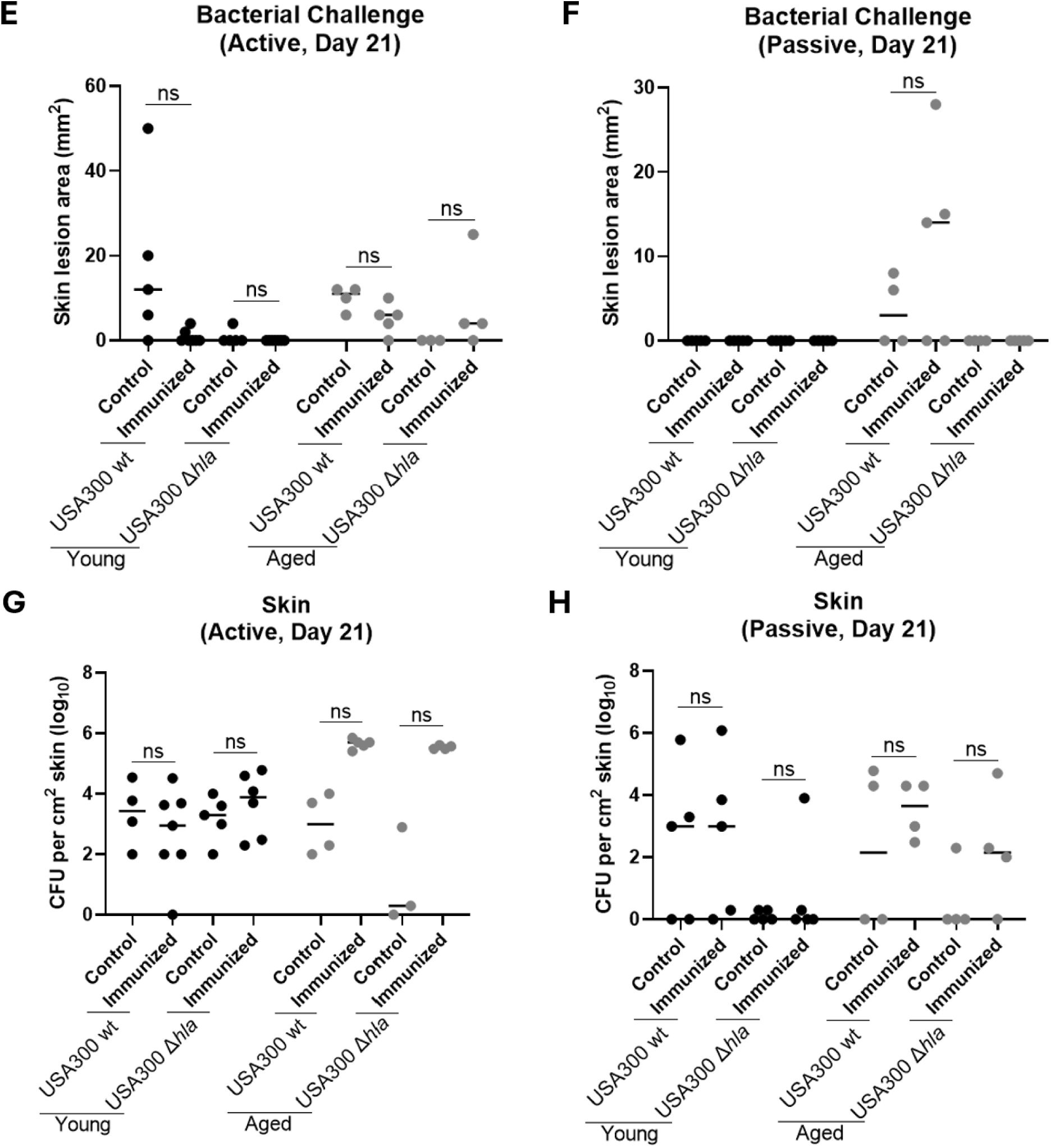
Quantitation of bacterial burden and skin lesions following anti-Hla immunization and Hla or *S. aureus* challenge of young and aged mice. Young and aged mice were administered active or passive anti-Hla immunization then challenged subcutaneously with Hla or WT or Δ*hla S. aureus*. (A) Active immunization - Skin lesion sizes 3 days after Hla challenge (n=5-9 mice). (B) Passive immunization - Skin lesion sizes 3 days after Hla challenge (n=5-9). (C) Active immunization - Skin lesion sizes 3 days after *S. aureus* infection (n=9-11 young, 4-7 aged). (D) Passive immunization - Skin lesion sizes 3 days after *S. aureus* infection (n=9-11 young, 4-7 aged). (E) Active immunization - Skin lesion sizes 21 days after *S. aureus* challenge (n=4-6). (F) Passive immunization - Skin lesion sizes 21 days after *S. aureus* challenge (n=4-5). (G) Active immunization - Bacterial burden 21 days post infection (n=4-6). (H) Passive immunization - Skin bacterial burden 21 days post infection (n=4-5). Each data point represents an individual mouse. Line represents mean. Kruskal-Wallis non-parametric one-way ANOVA test (A-H).

